# Towards pharmacokinetic profile predictions for monoclonal antibodies using sequence based machine learning derived parameters and compartmental modeling

**DOI:** 10.1101/2025.05.22.655474

**Authors:** Felix Jost, Henrik Cordes

## Abstract

This study presents a novel approach for predicting complete pharmacokinetic (PK) profiles of monoclonal antibodies (mAbs) based solely on their amino acid sequences, addressing a critical gap in early-stage therapeutic antibody development. While current methodologies rely on *in vivo* testing in specialized models like Tg32 mice or cynomolgus monkeys, our approach enables PK parameter prediction prior to molecule synthesis. Using a dataset of 118 diverse mAbs, we developed a physics-informed neural network (PINN) incorporating a two-compartment PK model. The four parameters of the PK model are predicted by a NN in which the numerical representation of the light and heavy chain sequences derived through sequence-attention and protein language model features serve as input. The model successfully predicted clearance and volume of distribution at steady state with remarkable accuracy, achieving predictions within 2-fold error for 10/13 and 13/13 mAbs in the test set, respectively. Concentration-time profiles were well-characterized with a median geometric mean fold error (GMFE) close to 2, with nearly 70% of predictions having GMFE less than 2. This proof-of-concept study demonstrates the feasibility of sequence-based PK prediction for mAbs, potentially reducing animal testing requirements while accelerating candidate selection in therapeutic antibody development. The approach aligns with regulatory initiatives promoting new approach methodologies and complements existing sequence-based predictions of protein structure and target binding during the design phase.

## INTRODUCTION

Monoclonal antibodies (mAbs) are a therapeutic modality providing potent and targeted (high specificity and target selectivity) medicine known for its less frequent administrations and off-target liabilities compared to small molecules ^1,2^. A clinical candidate should provide sufficient binding to its target in combination with favorable pharmacokinetic (PK) characteristics.

The potency of mAbs is typically assessed through *in vitro* binding assays, with subsequent validation in *in vivo* disease models. This dual approach facilitates the establishment of an *in vitro/in vivo* potency correlation, providing a comprehensive evaluation of the antibody’s therapeutic efficacy. With respect to PK, mAbs present distinct challenges compared to small molecules. While small molecule PK properties can be effectively translated from *in vitro* to *in vivo* systems, the analysis and extrapolation of *in vitro* PK characteristics to *in vivo* behavior for mAbs remains considerably more complex and challenging ^3^. An established strategy for human PK prediction is the determination of preclinical *in vivo* PK in Tg32 mice or cynomolgus monkeys (cyno) and its translation to human via allometric scaling achieving a human linear PK prediction within a 2-fold error ^4^. This approach holds true for mAbs with linear PK characteristics.

The elimination of mAbs within the context of PK characteristics encompass both excretion and catabolism. These elimination mechanisms can be further categorized into two primary pathways: nonspecific clearance and target mediated drug disposition (TMDD). The latter involves intracellular catabolism via lysosomal degradation following receptor-mediated endocytosis of the mAb-target complex ^5^. TMDD depends on target and biology and thus its impact varies across mAbs ^6^. In the present investigation, we exclusively examine linear PK as the characterization of TMDD necessitates multiple dose groups in PK studies. This constraint is not accommodated within our experimental design.

As outlined, the evaluation of PK characteristics relies on *in vivo* testing of clinical candidates at a certain stage of the screening cascade. *In vivo* experimentation for biological molecules presents significant ethical considerations and resource demands, surpassing those associated with small molecule studies. This increased complexity is primarily due to the necessity of utilizing specialized transgenic models, such as Tg32 mice, or non-rodent species, particularly cynos. Furthermore, the development of robust bioanalytical methods for each unique biological entity adds a substantial layer of complexity and resource utilization to the experimental process.

Different reasons still can lead to unfavorable PK properties ^3^. The primary objective in biological drug development is the identification of clinical candidates exhibiting optimal potency and PK properties. The volume of distribution at steady state (Vss) and clearance (CL) represent the fundamental PK parameters governing systemic exposure. While Vss demonstrates interspecies consistency due to limited tissue distribution kinetics and extent of monoclonal antibody (mAb) disposition, CL can exhibit considerable variability. This variability in CL is attributed to multiple antibody-specific properties, including atypical distribution patterns, non-specific binding phenomena, immunogenicity, molecular charge characteristics, isoelectric point, hydrophobicity, aggregation propensity, thermal stability, solution viscosity, and product purity. These physicochemical and biological attributes are systematically evaluated through a comprehensive panel of *in vitro* analytical assessments ^7–9^. In contrast, i*n vivo* testing is no high throughput screening such that *in vitro / in vivo* correlation (IVIVC) efforts together with *in silico* approaches have been developed to make *in vivo* CL predictions based on *in vitro* assay outcomes ^7^. Within this methodological framework, candidate molecules must be generated and undergo systematic evaluation through a comprehensive *in vitro* screening cascade during the lead identification phase before an IVIV extrapolation can be performed. These methodological approaches align with the recently published roadmap by the U.S. Food and Drug Administration that outlines a strategy for implementing the 3Rs principles (reduction, refinement, and replacement) of animal testing in preclinical safety evaluation. This initiative promotes the adoption of scientifically validated new approach methodologies, including microphysiological systems (organ-on-a-chip), computational toxicology models, and advanced *in vitro* assay platforms that recapitulate key aspects of human physiology and toxicological responses ^10^.

In addition to FDA-engaged initiatives on IVIVC and artificial intelligence / machine learning (AI/ML) methodologies that leverage *in vitro* data to predict individual PK parameters, our study presents a proof-of-concept approach wherein complete PK profiles of mAbs are predicted solely from their light and heavy chain amino acid sequences. This novel methodology aims to enable early CL, Vss, and PK profile predictions prior to molecule synthesis or *in vitro* data generation for early compound ranking and selection complementing the sequence based prediction of protein structure and target binding during design phase.

Similar to approaches for small molecules predicting PK profiles based on chemical structures, the sequence of mAbs is represented as a numerical vector which serves as input for a neural network architecture. Based on our findings for small molecules ^11^, we make use of a physics-informed neural network (PINN) in which a neural network is trained on concentration time profiles which are described by a two compartment model parametrized by the parameters V1, V2, CL, CL2. Betts et. al showed that for linear PK of mAbs a two compartment model adequately describes the biphasic linear PK profiles ^4^.

For the numerical representation of the mAbs, the python package Catpred is used ^12^. Catpred was developed to predict kcat, kinac, for substrates to certain enzymes. In this predictive methodology, amino acid sequences—specifically the light and heavy chains of mAbs in our application, as opposed to enzymes in the original approach—are converted into numerical representations through one to three distinct transformation methods. These transformations are subsequently concatenated to generate the final numerical representation of the antibody structure.

To the best of our knowledge, the prediction of complete PK profiles based solely on amino acid sequence has not yet been demonstrated in the literature. Recently, Genentech researchers reported the development of a binary classification system for distinguishing between high and low CL in cyno using sequence information with a dataset of comparable magnitude to our own. Their investigation emphasized the significant value of early CL prediction derived from sequence data and, in their future directions, proposed extending their methodology beyond binary classification to quantitative CL value estimation and comprehensive PK profile prediction ^8^.

## METHODS

### Numerical representation of amino acid sequence

Each mAb is represented as a vector of pooled features on the amino acid sequence level using sequence-attention and protein language model features. The light and heavy chains of a mAb are separately transformed into pooled features resulting in a concatenated numerical representation of size 1316+1316 = 2632. The python package CatPred ^12^ was used to calculate the numerical representations of mAb sequences which serve as input for the secondary machine learning method to predict the PK parameters. Therein, for the derivation of sequence attention (Seq-Attn) features, amino acid sequences were numerically encoded utilizing rotary positional embeddings analogous to those employed in the encoding layer of the ESM-2 pretrained language model (pLM). These encoded representations were subsequently processed through self-attention layers to capture long-range dependencies and contextual relationships across the sequences ^12^. Complementary to this approach, pLM features were extracted using the ESM-2 model (650 million parameters), which was pretrained on the UniRef50 dataset to capture evolutionary information embedded within protein sequences (see ^12^ for more information).

### Compartmental modeling

For each mAb a two compartment PK model with linear clearance from the central compartment was fitted to the mean concentration time profiles.

The fitted PK models and its estimated PK parameters are used for the calculation of the performance metrices introduced in the next section.

### Performance metrices

We utilized a summary of common PK metrics (C0, AUC, Cmin), percentage of predicted *in vivo* CL and Vss predictions within 2- and 3-fold error and geometric mean fold error (GMFE) between estimated and predicted concentration vs. time data.

### Physical informed neural networks

A PINN was set up using a two-compartmental PK model. As loss function, the logarithmic difference between PK concentration-time observations and predictions divided by the number of samples and the number of individuals in the batch was used. During model development, the number of hidden layers, the number of nodes, different activation functions and the batch size for the minibatch training including different learning rates were optimized.

### Data set generation & preparation

A comprehensive PK dataset was assembled, comprising 118 conventional monoclonal antibodies with 108 IgG1, 2 IgG2, 7 IgG4 and 1 mulgG2a-LALA isotypes evaluated during preclinical investigations conducted at Sanofi.

### Splitting strategy

The prepared data set was divided into training, validation, and test sets with a split of 80%, 10%, 10% using a random splitting strategy.

### Software

Compartmental modeling, descriptive statistics and visualization was performed within RStudio (version 4.2.0). For parameter estimation, the nonlinear least squares (nls) algorithm interfaced in nlmixr2 (version 2.0.8) was used.

The PINNs were set up in Julia^13^ (v1.10.4) with Flux^14^ (v0.14.19) and the native Vern7() ODE solver from the OrdinaryDiffEq^15^ (v6.89.0).

For the calculation of the numerical representation of the light and heavy chain of the mAb Catpred^12^ was used.

## RESULTS

### Data set

The dataset with 118 mAbs encompasses a diverse spectrum of PK characteristics especially non-specific clearance values quantified in transgenic Tg32 mice (1.51 – 88.7 mL/day/kg or 0.0629-3.69 mL/h/kg) and mean, median, minimal and maximal values for V1 (55.59 mL/kg, 47.32 mL/kg, [26.29-372.7]), V2 (85.05 mL/kg, 70.48 mL/kg, [17.65-644.8]) and CL2 (4.79, 3.39, [0.450-35.1]) from compartmental modeling. Figure S1 indicates that all concentration time profiles are adequately described by fitted two compartment models. Taking the threshold of 0.32 mL/h/kg or 7.68 mL/day/kg, defined as a reasonable threshold value for acceptable CL in Tg32^7,16,17^ and cyno^8^ the dataset consists of a slightly unbalanced distribution with 59% of mAbs having a slow CL lower than 0.32 mL/h/kg and 41% having a high CL higher than 0.32 mL/h/kg.

### Physics informed neural network

The best performing CMT-PINN was achieved with three full connected hidden layers with size of 128, *relu* activation function, *softplus* output activation function, batch size 32, epochs 4000 and the default learning rate 0.001 of Adam optimizer.

Next, we evaluated the prediction performance of the trained PINN on the test set comprising of 13 mAb PK profiles.

The PINN with the two compartment PK model adequately describes the mean concentration time profiles for each of the monoclonal antibodies in the test set for observed vs predicted concentrations. Figure 2 acknowledges this evaluation with a solid goodness of fit plot with data points around the line of identity stratified into the training, validation and test set.

The developed PINN predicts the *in vivo* clearance and the volume of distribution of 10 and 13 out of 13 mAb within a 2-fold error for one specific random data split, respectively. The accuracy to predict CL and Vss is visualized in Figure 3.

**Figure 1:**
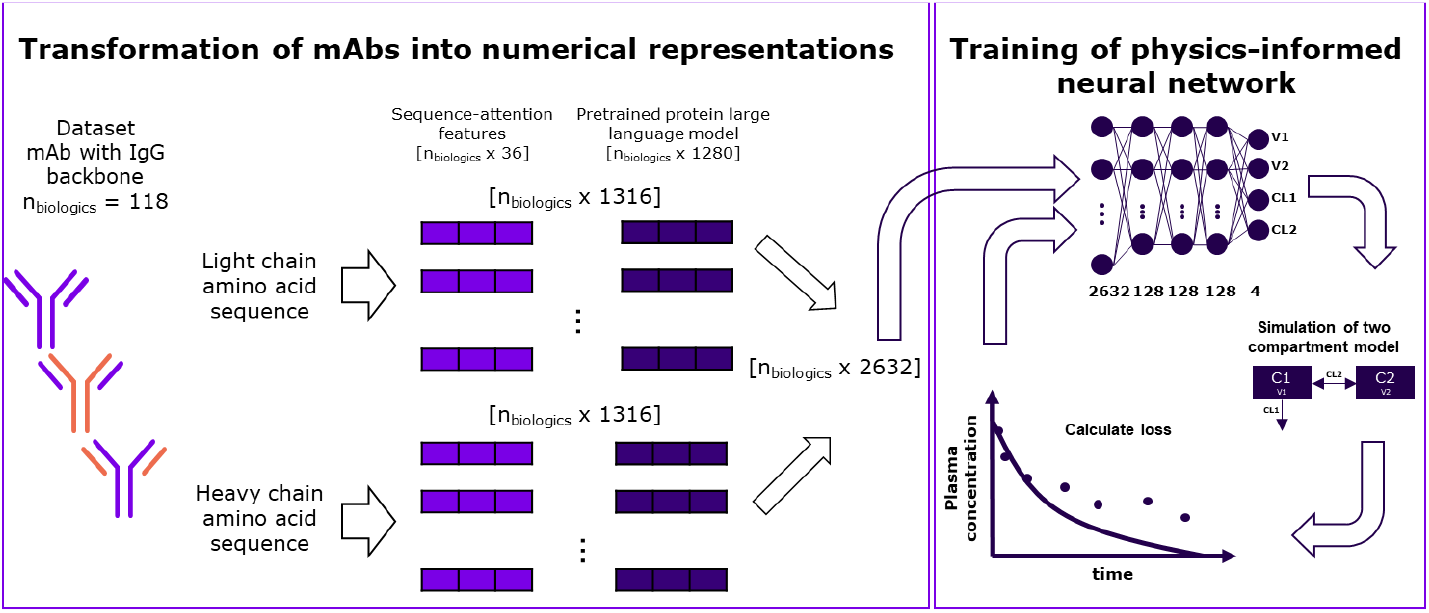
Schematic presentation of deep learning architecture for numerical representation of amino acid sequence and physics-informed neural network framework.

**Figure 2:**
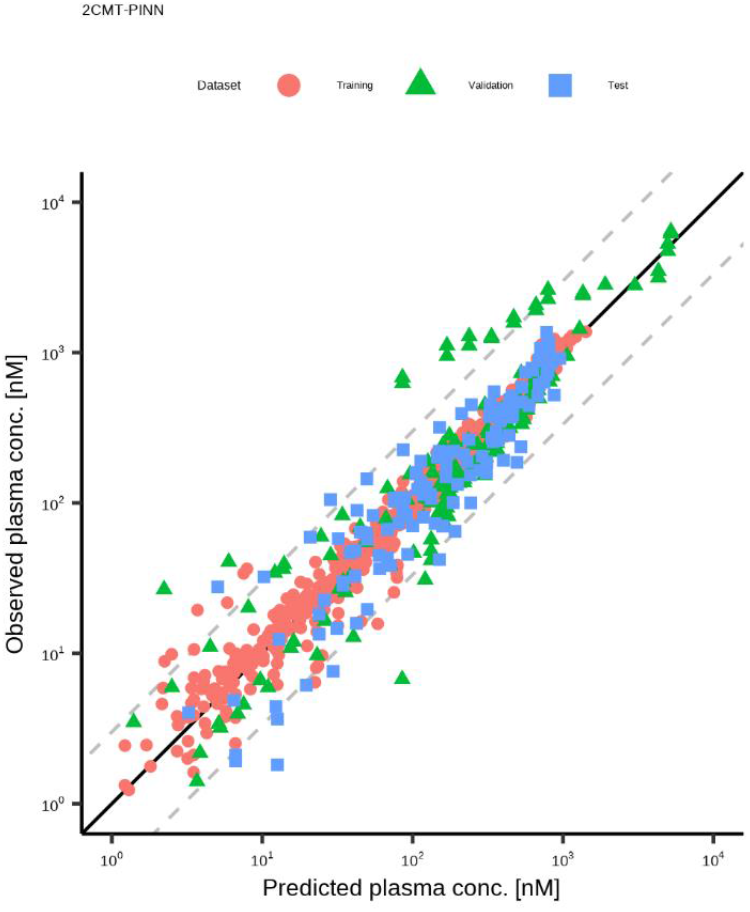
Predicted vs observed concentration data points separated into training, validation and test data sets.

**Figure 3:**
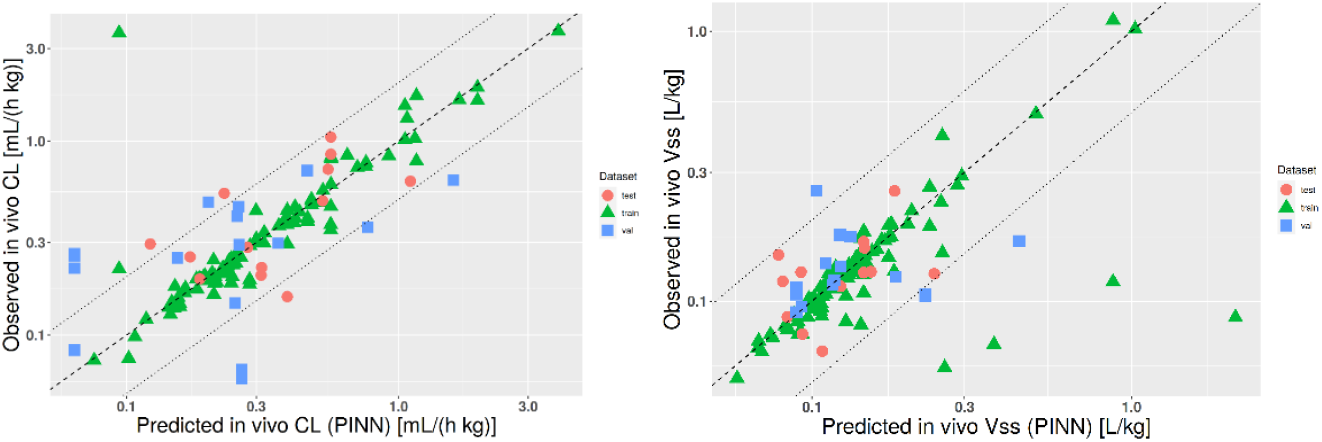
Predicted vs observed in vivo CL and Vss separated into training, validation and test data sets.

For the specific PK metrices, median relative errors for AUC, C0 and Cmin (at t=28 day) of 0.88, 1.06 and 0.53 reveal a slight underprediction of AUC and a 2-fold underprediction of Cmin (see Table 1).

**Table 1:**
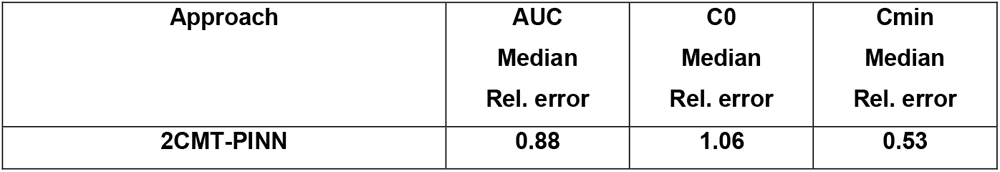
Median of relative errors between simulations and predictions for AUC, C0 and Cmin for the test data set.

GMFEs on simulated vs. predicted concentration time profiles result in a median GMFE value close to 2, nearly 70% of predictions with GMFE < 2 and only 7.7% of predictions with GMFE > 3 (see Table 2).

**Table 2:** Median and percentage of geometric mean fold error (GMFE) lower than 2,3 and 5 between simulated and predicted concentration data.

| Test-Set | 2CMT-PINN |
| --- | --- |
| Median GMFE | 1.88 |
| % GMFE < 2 | 69.2 |
| % GMFE < 3 | 92.3 |
| % GMFE < 5 | 100 |

## DISCUSSION and CONCLUSION

AI/ML approaches are heavily used in the design of therapeutic antibodies for sequence based predictions of structure and functions including affinity and potency ^18–20^. Although the translation of preclinical linear *in vivo* PK to human PK is well understood, the early prediction of *in vivo* PK characteristics purely based on amino acid sequence before compound generation is still challenging.

Current research efforts are focused on leveraging early-stage *in vitro* screening data for the prediction of *in vivo* PK characteristics through IVIVC methodologies ^7^. This approach holds validity and demonstrates utility at the developmental stage when candidate compounds have been synthesized and are amenable to *in vitro* testing.

In the present investigation, we advance the predictive paradigm to an earlier stage in the screening cascade, preceding actual molecule synthesis. Utilizing solely amino acid sequence information, we have developed a methodology for predicting *in vivo* CL, Vss and comprehensive PK profiles in Tg32 mice. This approach facilitates candidate ranking and enables preliminary human linear PK projections prior to physical compound generation, thereby potentially enhancing the efficiency of therapeutic antibody development ^4,9^.

In this study, we present a novel *in vivo* PK prediction framework that generates two-compartmental model parameters for transgenic Tg32 mice based solely on amino acid sequence information of mAbs. To our knowledge, this represents the first investigation demonstrating the prediction of complete PK profiles for mAbs utilizing sequence information alone. The methodology employs sequence attention and a pre-trained protein language model for numerical sequence representation, coupled with a PINN for PK parameter prediction. Using a specific randomized data partition, the model demonstrates promising predictive accuracy for both PK metrics and parameters, successfully reconstructing comprehensive concentration-time profiles.

The dataset utilized in this study encompasses 118 mAbs exhibiting substantial PK variability with CL values approximately 20% higher compared to the reported CL by Betts et al. for clinical mAb candidates ^4^. This diverse dataset enhances model training and generalization capabilities. The observed parameter estimates underscore the considerable PK variability that can manifest in mAbs that have not undergone optimization for clinical development. While acknowledging the general PK properties and variability typically associated with clinical mAbs, our dataset confirms an even higher degree of PK variability in non-optimized mAbs. This observation emphasizes the critical importance of identifying clinical candidates with acceptable PK properties during the screening cascade, as PK characteristics can significantly impact the developability and potential efficacy of therapeutic antibodies.

This proof-of-concept investigation establishes a foundation for further methodological refinements, which are systematically addressed in the subsequent sections. The preliminary findings presented herein identify several avenues for potential enhancement of the predictive framework.

The methodology described herein may be extrapolated to other biologic modalities, including nanobodies, hybrid constructs, and multispecific antibodies, contingent upon the availability of comparably sized datasets for each distinct modality. This approach ensures modality-specific training and validation of predictive models.

An alternative strategy worthy of exploration involves the integrated analysis of diverse large molecule modalities to identify sequence fragments that exert significant influence on PK behavior across modalities. These identified fragments could potentially serve as selective descriptors for PK predictions, facilitating a more universal approach to *in silico* PK modeling of biotherapeutics. This cross-modality analysis may elucidate common structural determinants of PK properties, potentially enhancing our understanding of structure-PK relationships in complex biologics.

Another field of investigation is the deep neural network architecture. The prediction approach uses a pretrained general large language model for the calculation of the numerical representation of mAbs. Mazrooei et. al ^8^ uses a pLM which was trained on mAb sequences and might be more specialized regarding feature calculation as the general pLM model from Catpred ^12^. In this context a dimension reduction on the descriptor level should be performed to reduce the possibility of overfitting. The current NN architecture has an input size of 2763 whereas the approach of Mazrooei et. al^8^ ended up with 15 features as input after permutation feature importance.

In this proof-of-concept investigation, we employed random data partitioning for model development and validation. Given the relatively limited dataset dimensions and the implementation of deep learning methodologies, more robust validation strategies may be beneficial in future iterations. Specifically, cross-validation techniques such as leave-one-out validation or Monte Carlo bootstrapping approaches could potentially enhance predictive accuracy and model generalizability. These advanced validation methodologies would provide more comprehensive assessment of model performance across different data subsets, thereby increasing confidence in the predictive capabilities of the framework when applied to novel molecular entities.

Lastly, the current prediction model was developed and validated using linear PK profiles, thereby not accounting for nonlinear PK such as TMDD. and neglects the aspect of nonlinear PK properties like TMDD. To address this limitation and expand the model’s applicability, future investigations should consider incorporating datasets with multiple dose groups for each mAb, enabling the characterization of individual TMDD profiles. Alternatively, the implementation of more mechanistically sophisticated mathematical frameworks, such as physiologically-based PK models tailored for biologics, could provide a more comprehensive representation of the complex PK behaviors observed in therapeutic antibodies. These enhancements would significantly augment the model’s predictive capacity and broaden its utility across diverse PK scenarios encountered in antibody therapeutics.

In conclusion, our findings represent a significant advancement toward establishing an AI/ML-driven research screening paradigm for biologic therapeutics. This work demonstrates a substantial expansion of sequence-based predictive methodologies beyond applications in structural and functional characterization, extending into the realm of comprehensive PK profile prediction. Such capabilities have the potential to fundamentally transform early-stage candidate selection and optimization processes in biotherapeutic development.

## ACKNOWLEDGEMENTS

*We would like to thank all Sanofi colleagues who developed the compounds and generated the in vivo PK data for this analysis*.

## AUTHOR CONTRIBUTIONS

F.J. and H.C wrote the manuscript; F.J. designed and performed the research; F.J. and H.C analyzed the data.

## SUPPLEMENTARY

**Figure S1:**
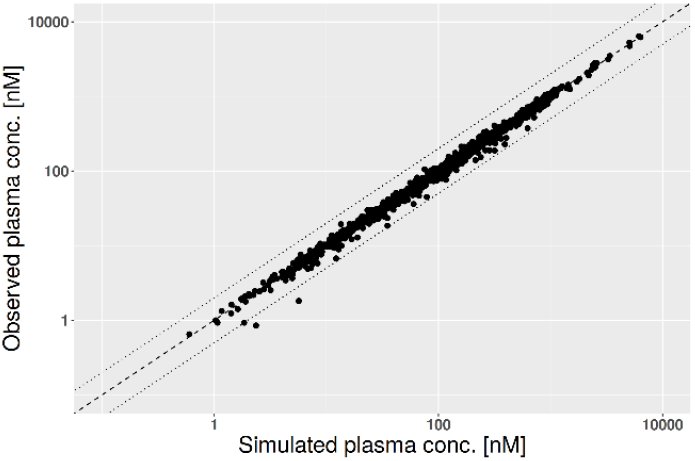
Simulated vs observed concentration data points after parameter estimation.

